# Intestinal Mucin Is a Chaperone of Multivalent Copper

**DOI:** 10.1101/2022.01.02.474741

**Authors:** Nava Reznik, Annastassia D. Gallo, Katherine W. Rush, Gabriel Javitt, Yael Fridmann-Sirkis, Tal Ilani, Noa A. Nairner, Kelly N. Chacón, Katherine J. Franz, Deborah Fass

## Abstract

Mucus protects the body by many mechanisms, but a role in managing toxic transition metals was not previously known. Here we report that secreted mucins, the major mucus glycoproteins coating the respiratory and intestinal epithelia, are specific copper-binding proteins. Most remarkably, the intestinal mucin, MUC2, has two juxtaposed copper binding sites, one that accommodates Cu^2+^ and the other Cu^1+^, which can be formed *in situ* by reduction with vitamin C. Copper is an essential trace metal because it is a cofactor for a variety of enzymes catalyzing electron transfer reactions, but copper damages macromolecules when unregulated. We observed that MUC2 protects against copper toxicity while permitting nutritional uptake into cells. These findings introduce mucins, produced in massive quantities to guard extensive mucosal surfaces, as extracellular copper chaperones and potentially important players in physiological copper homeostasis.

## INTRODUCTION

Giant mucin glycoproteins are secreted from cells to form protective mucus hydrogels covering the exposed surfaces of internal organs such as the lung and intestine (McGuckin et al., 2011; Benam et al., 2018). These hydrogels guard the underlying epithelium from pathogens and other hazardous matter entering from the outside world, while permitting nutrient absorption and gas exchange. Mucin gels store antimicrobial molecules that participate in innate immunity (Padra et al., 2019; Hoffmann, 2021). Mucin glycoproteins also house and feed the microbiome, lubricate tissue surfaces, and may facilitate the removal of contaminants and waste products from the body (Paone & Cani, 2020). These numerous activities are enabled by the large sizes, dynamic conformations, and extensive intermolecular interactions of mucins. However, the same features that contribute to mucin functionality also complicate the study of these glycoproteins. Consequently, many questions remain regarding the mechanisms of mucin assembly, their physical and chemical capabilities, and their physiological contributions to the critical interfaces between animals and their environments.

We recently determined, using cryo-electron microscopy (cryo-EM), the structure of the amino-terminal region of the intestinal mucin MUC2 in the context of a filament proposed to represent a bioassembly intermediate (Javitt et al., 2020). We noticed in the VWD1 (D1) region of this structure (Figures 1A and 1B) an intriguing congregation of histidine and methionine amino acids, including Met146, Met154, His277, His324, and Met 326, arranged with the side chains pointing toward one another. Most of these amino acids are highly conserved in vertebrate MUC2 orthologs (Figure 1C). Inspection of human mucin paralogs revealed that the histidines are found in other gel-forming mucins (*i.e*., MUC5AC and MUC5B), but the methionines are largely restricted to MUC2 (Figure 1D). In the genetic code, histidine is represented by two codons and methionine by only one, so these amino acids could readily have been replaced in evolution if they were not under strong functional selection.

**Figure 1.**
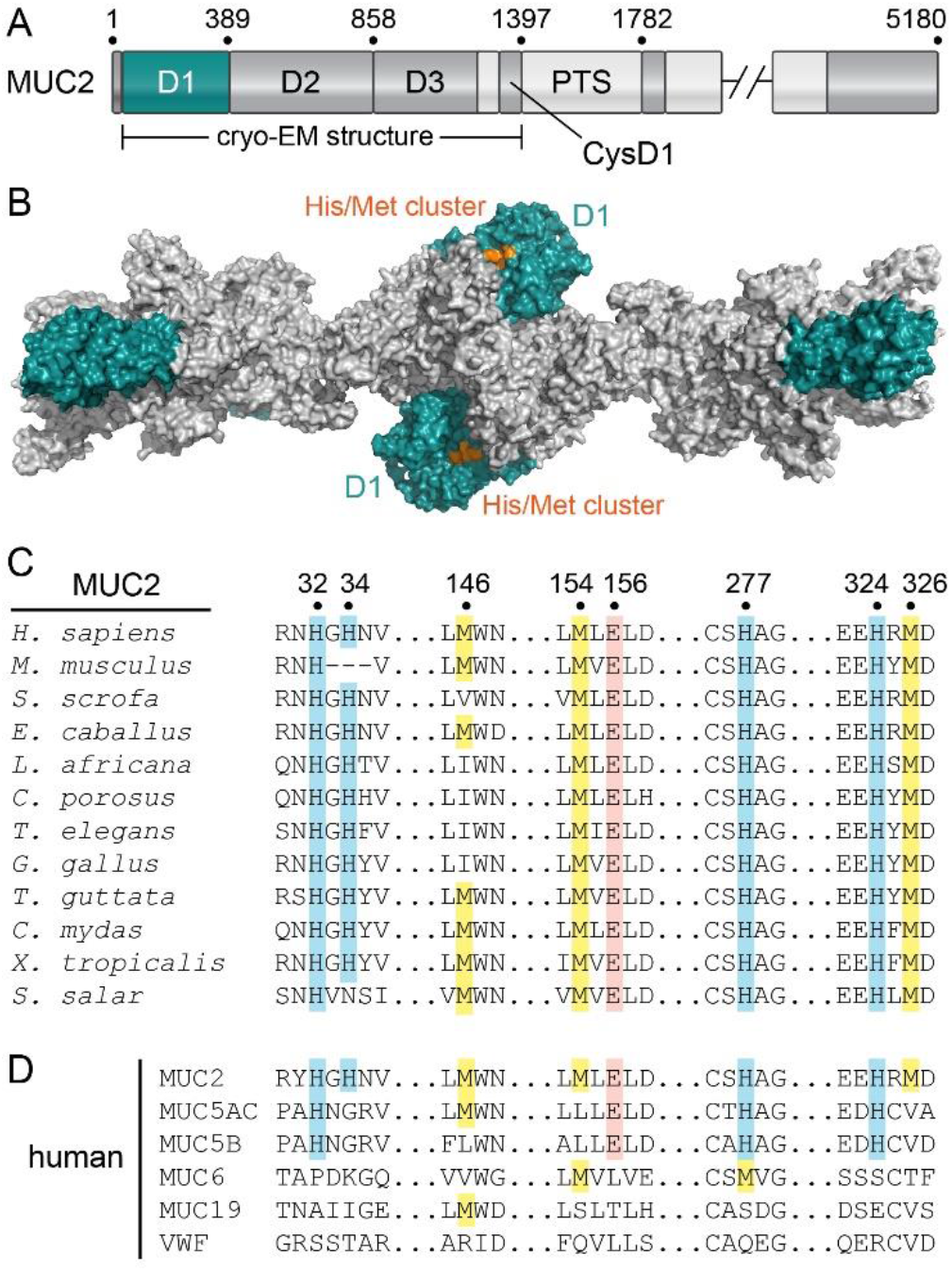
Histidines are Conserved in Intestinal and Respiratory Mucins, Methionines Only in Intestinal Mucins. (A) Domain map of human MUC2. The VWD1 (D1) domain is colored teal. “PTS” refers to low-complexity segments rich in proline and O-glycosylated serine and threonine. Numbers indicate amino acid positions at key domain boundaries in the MUC2 precursor (UniProt Q02817). (B) Filament formed by the amino-terminal region of MUC2 (Javitt et al., 2020), shown as a molecular surface representation. Orange indicates a cluster of histidines and methionines. (C) Segments of an amino acid sequence alignment of MUC2 orthologs from representative animal species. Histidines at the indicated positions are highlighted in blue, methionines in yellow, and a glutamic acid relevant to this work in pink. The human MUC2 sequence is taken from NCBI NP_002448.4. (D) Sequences of human gel-forming mucins and the related blood clotting protein von Willebrand factor (VWF). MUC5B and MUC5AC are gel-forming mucins in airway mucus (Benam et al., 2018).

Histidines and methionines are common copper ligands in proteins (Rubino & Franz, 2012). Copper is bound by proteins as a cofactor in a variety of important redox enzymes, to which is it delivered by dedicated transporters and chaperones in the circulation and in cells (Rubino & Franz, 2012; Linder, 2016; Magistrato et al., 2019). Dysregulation of copper handling due to genetic mutations impairs vascular, hepatic, and immune functions, and deficiency or mis-localization of copper is associated with the pathophysiology of neurodegenerative and other diseases (DiNicolantonio et al., 2018; Acevedo et al., 2019). Despite the importance of copper use and regulation in the body, little is known about the initial encounter with dietary copper or about copper management at mucosal surfaces. This gap is largely due to the heterogeneity of mucosal environments and, as noted above, limited knowledge about the complex macromolecules that function there. Here we present structural and biochemical evidence that mucins are specific copper-binding proteins. We show that the intestinal mucin engages copper in multiple oxidation states, and we demonstrate the functional benefits of copper regulation by mucin.

## RESULTS

### Gel-forming Mucins Bind Copper

To test whether MUC2 binds copper, the isolated D1 assembly was prepared, incubated with an equimolar amount of CuSO_4_, and crystallized (Table S1). The X-ray crystal structure revealed a Cu^2+^ binding site located at the interface of multiple subdomains near the protein surface (Figures 2A and S1A). Though histidines are also prevalent in zinc binding sites (Ireland & Martin, 2019), no evidence of zinc was seen in D1 crystals grown in the presence of ZnCl_2_ (data not shown). Microscale thermophoresis (MST) metal binding assays in solution supported a preference for Cu^2+^ (Figure S1B), and isothermal titration calorimetry (ITC) yielded an apparent logK_D_ of −13 for Cu^2+^ binding by D1 (Figure S1C and Table S2).

**Figure 2.**
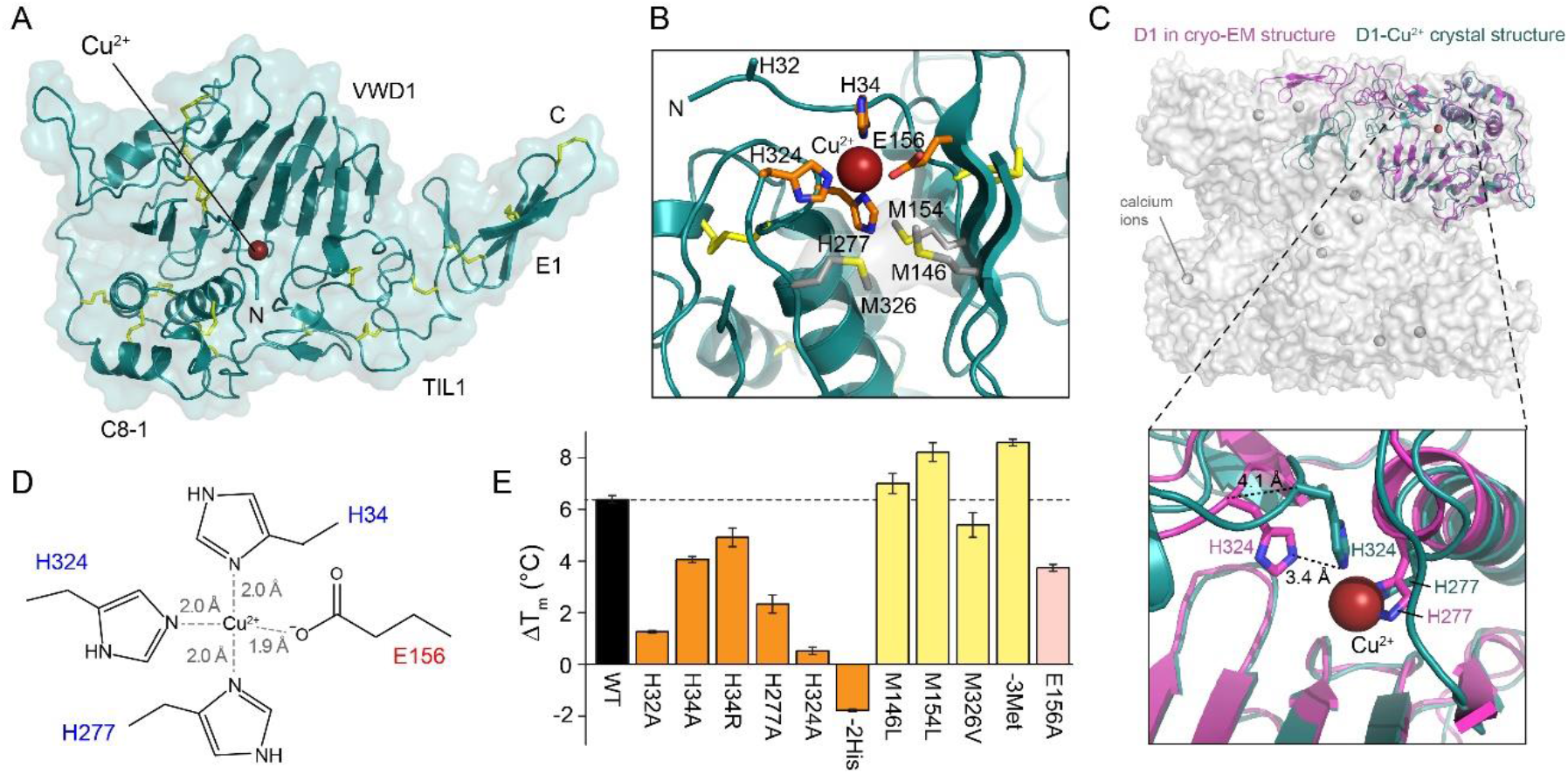
MUC2 Binds Cu^2+^. (A) MUC2 D1 Cu^2+^-bound crystal structure. A backbone cartoon is shown within a semi-transparent surface, and the dark red sphere is the copper ion. Disulfide bonds are yellow. Polypeptide termini are labeled “N” and “C.” Domains are labeled according to convention (Nilsson et al., 2014). See also Figures S1-S4. (B) Ligation of Cu^2+^ in MUC2 D1. Methionines (Met146, Met154, Met326) did not participate in Cu^2+^ coordination, and the Met sulfurs were more than 6 Å from the Cu^2+^. The His32 side chain was disordered in the crystal. (C) Comparison of the MUC2 D1 crystal structure with D1 in the cryo-EM structure of the filamentous assembly intermediate (Figure 1B). A cartoon representation of one D1 assembly is colored magenta within a semi-transparent surface representation of a unit in the filament (Figure 1B). The crystal structure of MUC2 D1 bound to Cu^2+^ is superposed in dark teal. The D1 assembly is rotated by about 180° compared to panel A. Calcium ions bound at specific acidic motifs found in mucins and the related von Willebrand factor are indicated (Dong et al., 2019; Javitt et al., 2019). The zoomed view below shows how movement of the TIL1 domain brings His324 closer for Cu^2+^ coordination. (D) Illustration of the Cu^2+^-coordination site with ligand-metal distances obtained from EXAFS. (E) Cu^2+^ binding increased the midpoint of thermal denaturation of MUC2 D1 by about 6 °C (black bar), measured using DSF at pH 5.7. The differences between the denaturation temperature with and without Cu^2+^ (∆T_m_) are shown for wild type (WT) and the indicated mutants, colored according to amino acid type mutated. His34 was mutated also to arginine because the UniProt entry Q02817 contains arginine at this position. The label “-2His” refers to a double H277A/H324A mutant. The label “-3Met” refers to a triple M146L/M154L/M326V mutant. Errors are standard deviation, n=3.

Cu^2+^ was coordinated in the MUC2 D1 crystal structure by three histidines and a glutamate (Figure 2B). In addition to His277 and His324, which drew closer together in the crystal structure compared to their positions in the cryo-EM cluster (Figure 2C), the amino-terminal region of D1 became ordered in the crystal, providing His34 for metal coordination. The highly conserved Glu156 (Figure 1C) completed the Cu^2+^ coordination sphere. The three methionines present in the cryo-EM cluster, Met146, Met154, and Met326, did not contribute to Cu^2+^ binding in the D1 structure and were positioned with their sulfur atoms about 6 Å away from the metal (Figure 2B). Extended X-ray absorption fine structure (EXAFS) of Cu^2+^-loaded MUC2 D1 was consistent with ligation to three multiple scattering histidine residues and one N/O ligand (Figure S2A and Table S3) at the indicated distances (Figure 2D). Though other proteins bind Cu^2+^ with similar ligands, the MUC2 geometry is distinct (Figure S3).

Supporting the structural studies, Cu^2+^ addition to MUC2 D1 increased the temperature midpoint of its thermal unfolding transition measured by differential scanning fluorimetry (DSF) (Figures 2E and S4A). This phenomenon was particularly dependent on His277 and His324 but independent of the methionines (Figure 2E). Interestingly, the effect of mutating His32, which is highly conserved (Figures 1C and 1D) but had a disordered side chain that did not participate in Cu^2+^ ligation in the crystal structure (Figure 2B), was similar in magnitude to mutation of the Cu^2+^ ligands His277 or His324 (Figure 2E). The impact of mutating His34, seen to bind Cu^2+^ in the crystal (Figure 2B), was more moderate (Figure 2E). It is possible that both His32 and His34 can engage in Cu^2+^ ligation, but crystal packing favored His34 participation. In light of the conservation of the histidines and glutamic acid in other major gel-forming mucins (Figure 1D), we hypothesized that these mucins also coordinate Cu^2+^. Indeed, D1 of the murine MUC5B ortholog (denoted Muc5b), a representative respiratory mucin, showed increased thermal stability in the presence of Cu^2+^ as measured by DSF (Figure S4B), and MST confirmed Muc5b Cu^2+^-binding activity (Figure S4C).

### Reduced Copper is Captured at a Neighboring Site in MUC2

Though the MUC2 D1 methionines did not engage in Cu^2+^ coordination (Figure 2B), the proximity of these conserved amino acids to the Cu^2+^ binding site could not be overlooked. Methionines often participate in Cu^1+^ binding (Rubino & Franz, 2012), leading to the hypothesis that MUC2 has distinct sites for Cu^2+^ and Cu^1+^. Intriguingly, Cu^2+^ bound by D1 was reduced to Cu^1+^ by ascorbate (vitamin C) and by the dietary flavonoid quercetin (Figure 3A). This observation inspired the addition of ascorbate to the Cu^2+^-bound MUC2 D1 crystals. Diffraction data collected from these crystals revealed a strong peak of electron density between the methionines, while the density corresponding to the histidine-dominated Cu^2+^ site had disappeared (Figure 3B). The resolution of the diffraction data from the ascorbate-treated crystals was not sufficient to precisely measure sulfur-copper distances, but EXAFS of MUC2 D1 in solution confirmed Cu^1+^ binding and provided structural information on the Cu^1+^ coordination environment. Upon treatment of Cu^2+^-bound MUC2 D1 with ascorbate, the ligation changed dramatically from three-His one-N/O to three sulfur ligands with an average Cu–S distance of 2.317 Å and DW of 0.011 Å^2^ (Figure S2B and Table S3), which agrees well with data on other methionine Cu^1+^ protein systems (Chacón et al., 2014; Martin-Diaconescu et al., 2016). These changes in coordination environment and redox state were also indicated by the differences in the copper edges of the raw X-ray scattering data (Figure 3C).

**Figure 3.**
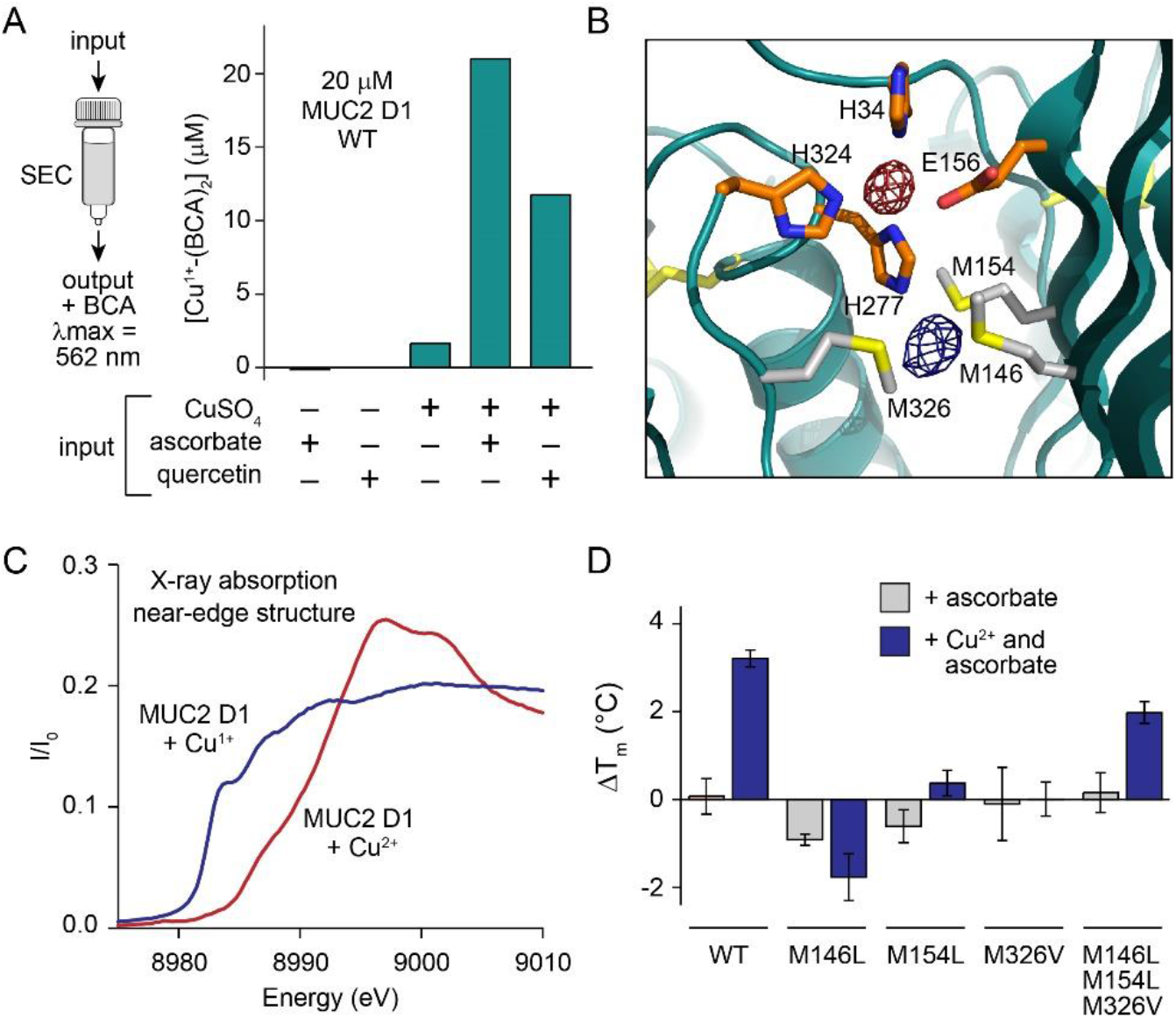
Reduced Copper is Transferred to a Nearby Methionine Cluster in MUC2. (A) The indicated additions were made to MUC2 D1. Buffer was rapidly exchanged on a size exclusion column (SEC) to remove excess reductant or unbound copper, and Cu^1+^ retained by 20 μM protein was detected using BCA. (B) Fo-Fc difference map density, calculated using the diffraction data from ascorbate-treated MUC2 D1-Cu^1+^ crystals phased using MUC2 D1 coordinates from which the Cu^2+^ ion had been removed. Map is displayed at 4.5σ (dark blue). For comparison, Fo-Fc map density of the Cu^2+^ crystal calculated using phases from the same model is superposed and displayed at 8σ. The ribbon diagram was made using the coordinates of the Cu^2+^-bound form without displaying the Cu^2+^. (C) Normalized X-ray absorption near-edge structure (XANES) spectrum of MUC2 D1 supplied with CuSO_4_ is shown in red. The blue spectrum was measured after treatment of the Cu^2+^-bound MUC2 D1 with ascorbate. The first inflection point of MUC2 D1 Cu^1+^ was calculated to occur at 8982.4 eV, while the first inflection point of MUC2 D1 Cu^2+^ was at 8986.0 eV. See also Figure S2. (D) The differences in the midpoints of the thermal unfolding transitions between untreated protein and protein after the indicated treatments are plotted. The methionines were required for the stabilization against thermal denaturation provided by Cu^1+^. It is possible that the triple mutant (M146L/M154L/M326V) retained some stabilization because the copper remained bound to the histidines and was not transferred to the defective Cu^1+^ site. Errors are standard deviation, n=3. See also Figures S4 and S5.

Mutagenesis further supported the importance of the methionines in Cu^1+^ binding. Cu^1+^ increased the temperature of the MUC2 D1 thermal unfolding transition to about half the extent as Cu^2+^, and its effect was entirely undermined by mutating Met146, Met154, or Met326 (Figure 3D). Furthermore, mutation of methionines lowered the affinity for Cu^1+^ as measured by competitive titrations (Figure S5). Muc5b, which naturally lacks the methionines, was not stabilized by Cu^1+^ (Figure S4D). Together, the experiments described thus far provide structural and biochemical evidence for a two-tiered copper binding environment in MUC2, in which histidines dominate the Cu^2+^ coordination and nearby methionines capture Cu^1+^.

### MUC2 Blocks ROS Formation and Protects Colon Cells from Copper Toxicity

As shown, Cu^2+^ bound to MUC2 D1 was reduced by ascorbate (Figure 3A). However, unlike copper in solution, which catalyzes the consumption of oxygen and depletion of the antioxidant (Buettner & Jurkiewicz, 1996), copper bound to MUC2 D1 did not engage in futile cycling of electrons (Figure 4A). This result indicates that MUC2 D1 stabilizes Cu^1+^ even in the presence of oxygen in a manner that prevents accumulation of reactive oxygen species.

**Figure 4.**
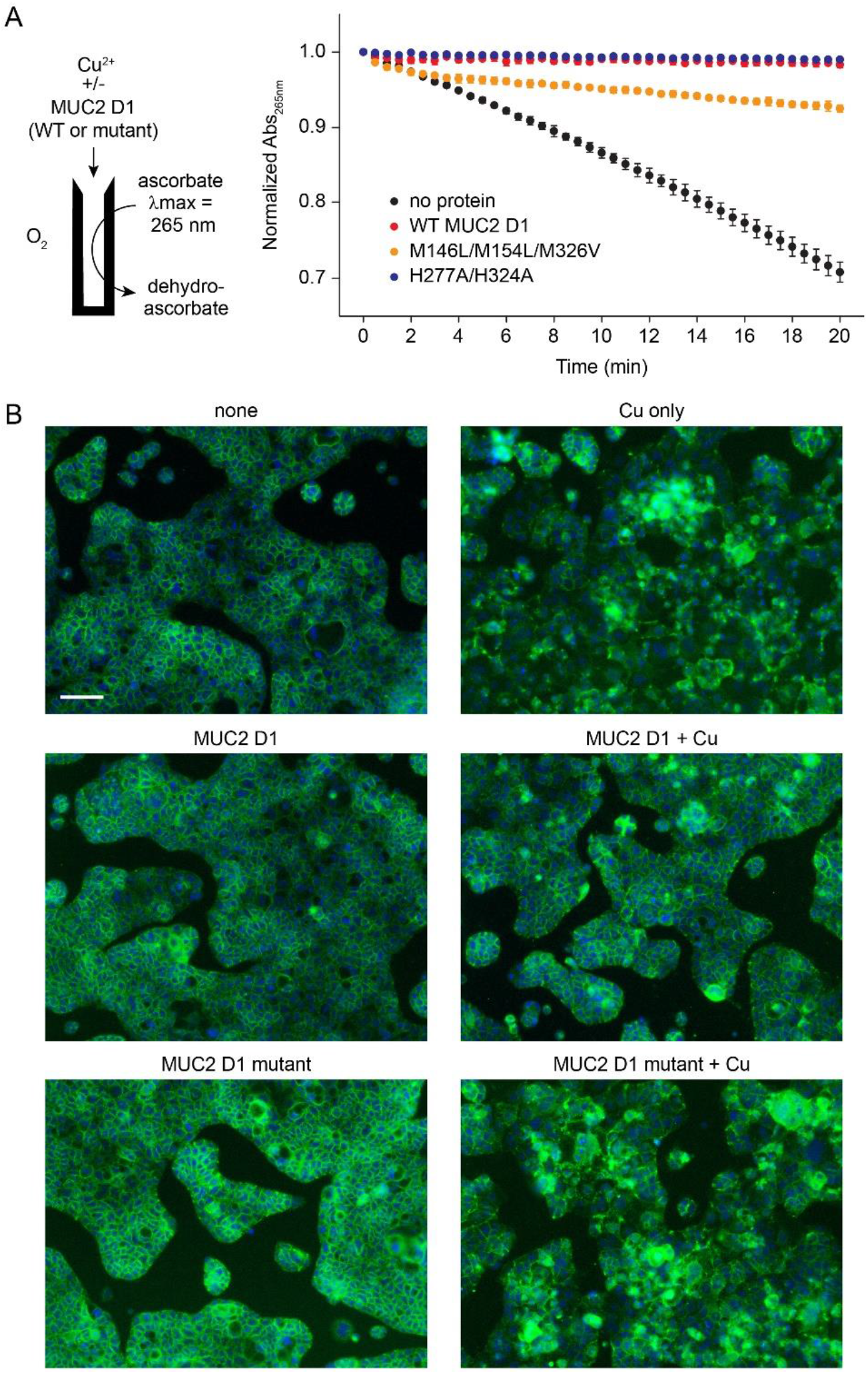
MUC2 D1 Prevents the Generation of Reactive Oxygen Species and Protects Colon Cells Against Toxic Copper Concentrations. (A) MUC2 D1 WT (red) and H277A/H324A (blue) prevented Cu^2+^-driven ascorbate oxidation, whereas the triple methionine mutant (M146L/M154L/M326V) (orange) decreased the rate of oxidation compared to Cu^2+^ alone (black). (B) Caco-2 colon cells were subjected to the indicated treatments in serum-free medium and then stained with DAPI (nuclei, blue) and labeled for E-cadherin (green). Cultures treated with a high copper concentration (250 μM) in the presence of equimolar wild-type MUC2 D1, but not of a H324A/M326V mutant, retained a normal appearance. Scale bar is 100 μm.

Based on this finding, we hypothesized that MUC2 might protect intestinal cells from toxic copper concentrations. E-cadherin labeling of Caco-2 human colorectal adenocarcinoma cells was used to report on the integrity of cell-cell contact sites and the health of the monolayer. Addition of 250 μM Cu^2+^ in serum-free medium resulted in dramatic loss of the organized networks of E-cadherin seen in control cultures (Figure 4B). Supplying 250 μM MUC2 D1 together with the 250 μM Cu^2+^ almost completely reversed this phenomenon, but a D1 mutant with perturbed copper binding sites, H324A/M326V, was unable to protect (Figure 4B).

### MUC2 Permits Copper Uptake into Cells

The ability of MUC2 D1 to neutralize the toxicity of excess copper led us to ask whether D1 completely sequesters extracellular copper, or rather allows bioavailable copper uptake for beneficial physiological processes. To address this question, we investigated whether nutritional levels of copper supplied with D1 can be taken up by copper-starved cells. Caco-2 cells were deprived of copper until a severe drop was seen in the level of the mitochondrial electron transport chain component cytochrome c oxidase subunit 1 (COX1), which requires copper for function (Getz et al., 2011). Limited amounts of copper, 0.1 or 1 μM, were then re-supplied, either alone or in the presence of stoichiometric or excess MUC2 D1, and the recovery of COX1 levels was monitored. Recovery proceeded even with a five-fold excess of D1 over copper, in a concentration range substantially over the dissociation constant, suggesting that D1 can release needed copper to cells (Figures 5A and S6). Hinting at a mechanism for Cu^1+^ transfer, the Cu^1+^ ligands in MUC2 D1, though buried within the protein, are adjacent to a small interior tunnel with two portals at the protein surface (Figure 5B). Together these findings show that MUC2 guards cells from the toxicity of excess copper without blocking supply of the low levels of copper required to maintain physiological processes.

**Figure 5.**
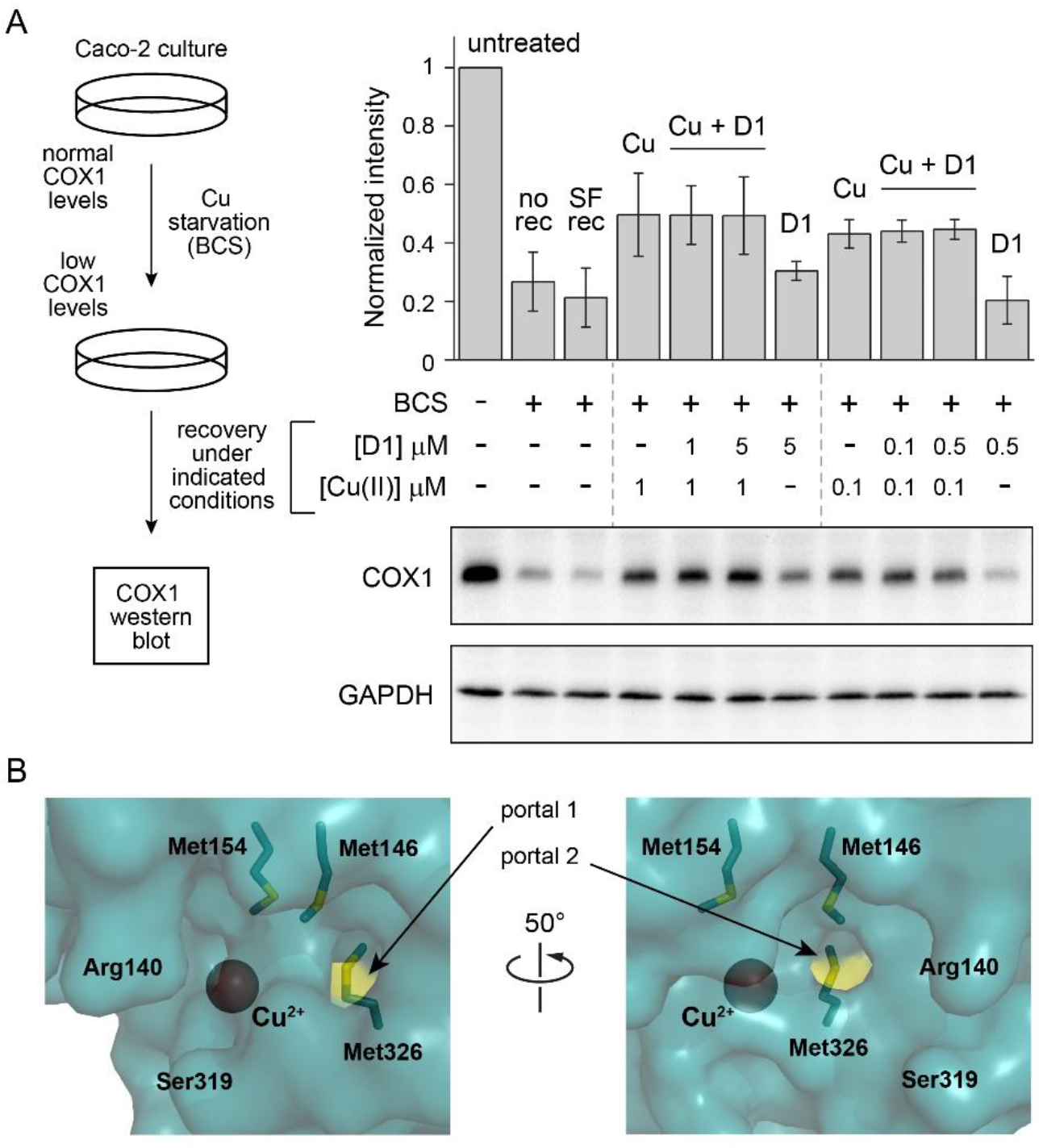
MUC2 D1 can release copper to cells. (A) Supplying copper bound to MUC2 D1 restored COX1 levels in copper-starved Caco-2 cells. “SF rec” indicates that only serum-free medium was supplied during the recovery period, whereas the “no rec” sample was harvested without recovery. Dashed gray lines separate sets of samples with higher and lower copper and D1 concentrations during recovery. See also Figure S6. (B) Met326 is exposed to the exterior of the protein through two small portals. A surface representation of the Cu^2+^-bound MUC2 D1 crystal structure is shown with sulfur atoms colored yellow.

## DISCUSSION

Mucins are multifunctional molecules that promote mucosal health by restricting access of pathogens to the epithelial cell surface, storing factors that participate in innate immunity, and managing the microbiome (McGuckin et al., 2011; Johansson & Hansson, 2016; Chairatana & Nolan, 2017; Benam et al., 2018; Bankole et al., 2021). Moreover, secreted mucins were hypothesized to act not only as a protective barrier but also as regulators of small molecule access to membrane-embedded receptors and transporters (Strous & Dekker, 1992). However, investigating the molecular mechanisms of mucin function has long been limited by the lack of high-resolution mucin structures. Remarkably, a specific role for mucins in managing copper was revealed when structural information became available to inspire and inform the experiments reported in this work.

The studies presented here show that mucin D1 assemblies are specific for Cu^2+^ over a variety of other metals tested. Weak binding to zinc was also detected, but no evidence could be obtained from crystallography that this ion binds at the Cu^2+^ site. The specialization of the mucin D1 assembly for copper binding contrasts with a separate, conserved calcium binding site present in all three D assemblies in the amino-terminal region of MUC2 (Javitt et al., 2020), as well in the three homologous D assemblies of the related blood clotting glycoprotein von Willebrand factor (Dong et al., 2019; Javitt et al., under revision). Calcium was not observed in the crystals of Cu^2+^-bound MUC2 D1, perhaps because it was stripped by the citrate present in the crystallization solution, though MUC2 D3 was also crystallized with citrate and retained its calcium (Javitt et al., 2019). All solution measurements of Cu^2+^ binding were likely performed on Ca^2+^-loaded MUC2 D1, as addition of Ca^2+^ had no effect on the protein in MST (Figure S1B) or DSF (Figure S4A) experiments.

Copper and calcium binding occur in different manners in MUC2. Ca^2+^ is bound locally within the VWD domains of the D assemblies, whereas Cu^2+^ is bound at the interface between three separate domains in D1: VWD1, C8-1, and TIL-1 (Figure 2A). One Cu^2+^ ligand is provided by each of these domains, and the fourth ligand is supplied by the flexible region at the amino terminus of the protein, just upstream of the VWD1 domain (Figure 2B). The Cu^1+^ ligands, in turn, are provided by the VWD1 and TIL-1 domains, without the participation of C8-1. The binding of copper between multiple domains suggests that affinity for copper is tunable by factors or events, such as interactions with other proteins, that may affect the relative orientations of those domains. Indeed, the TIL-1 and E-1 domains are flexible relative to the VWD1 and C8-1 domains, affecting the availability of the key copper ligand H324 (Figure 2C).

With detailed structural and biochemical information now available, a physiological role for copper binding by mucins can be considered. Just as mucins themselves have many functions, copper binding by mucins may serve various purposes. MUC2 is expressed at high levels in all parts of the intestine, including both aerobic and anaerobic environments and regions with different digestive, absorptive, and excretive roles (Audie et al., 1993; Paone & Cani, 2020). In addition to protecting the intestines and other mucosal surfaces against copper toxicity, copper binding by mucins may facilitate the conversion of dietary copper (largely Cu^2+^) to the form transported across the enterocyte membrane (thought to be Cu^1+^). The mechanisms by which dietary copper passes from the lumen of the digestive tract into enterocytes are debated (Zimnicka et al., 2007; Nose et al., 2010; Pierson et al., 2019). Hypothetically, MUC2 may interface structurally or functionally with the transmembrane copper transporter Ctr1, which uses methionines to coordinate Cu^1+^ in its pore (Ren et al., 2019), or with other proteins putatively involved in intestinal copper uptake (Nose et al., 2006). Mucins may also participate in nutritional immunity (Lopez & Skaar, 2018) by depriving pathogens of copper, and MUC2 may cooperate with bile (Linder, 2020) for the safe excretion of excess copper. These activities need not be mutually exclusive. Further exploration of these various possibilities will help integrate mucins into models of physiological copper management (Shanbhag et al., 2021).

It is well known that copper is carefully regulated in biology. Transfer of copper within eukaryotic cells and in the bacterial periplasm is done by direct protein-protein contacts and ligand exchange rather than by release of copper into solution (Robinson & Winge, 2010; Nevitt et al., 2012). The players and mechanisms of copper chaperoning inside cells and in blood are relatively well understood (Nevitt et al., 2012; Linder, 2016; Magistrato et al., 2019), but how the body manages copper before it reaches those sites has been obscure. The lung and intestinal mucosa are the largest, most important, and most vulnerable of exposed physiological surfaces, so it is reasonable that a mechanism would have evolved for restraining an essential but toxic element in these environments. The discovery that multiple gel-forming mucins are copper binding proteins, with an elaborate two-tiered site for distinct copper redox states in intestinal mucin (Figure 6), introduces intriguing new players into the field of copper regulation and utilization in the human body.

**Figure 6.**
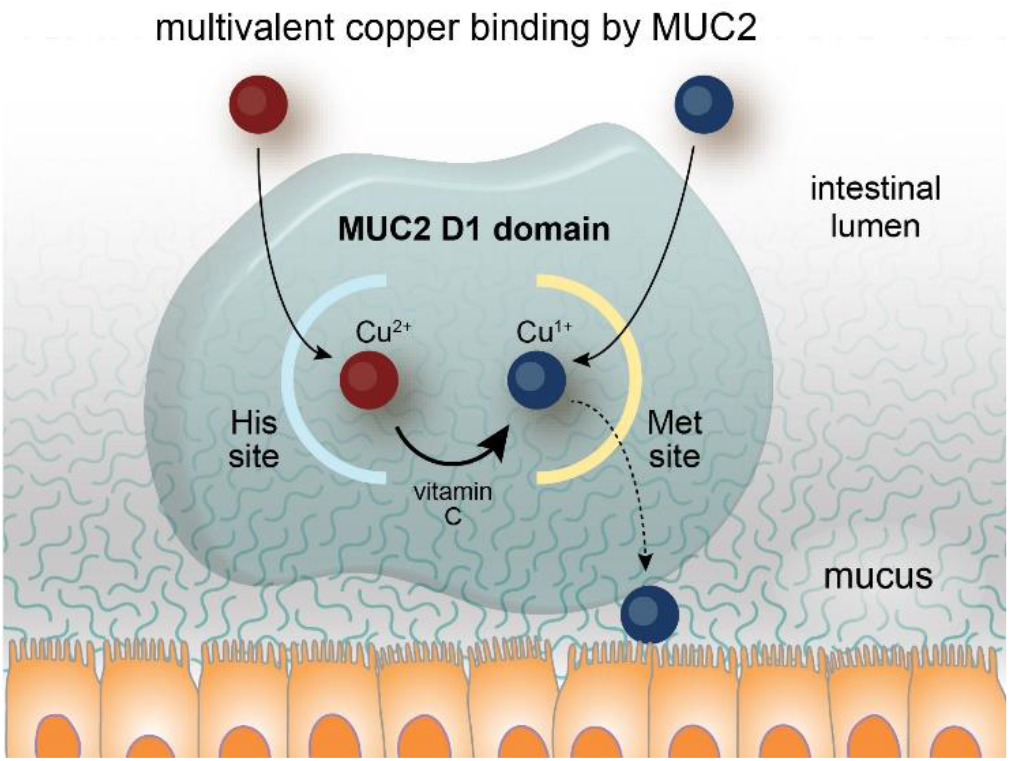
Summary of Copper Chaperoning by MUC2. A schematic illustration of the MUC2 two-tiered copper binding site, in which a histidine-rich site (His site) binds captures Cu^2+^ and a methionine-rich site (Met site) captures Cu^1+^. MUC2-bound Cu^2+^ can be reduced to Cu^1+^ by vitamin C (ascorbate) or other dietary antioxidants. Bound Cu^1+^ is protected by MUC2 from oxidation in aerobic environments, but copper, putatively in the form of Cu^1+^, can be released by MUC2 for nutritional delivery to cells (dashed arrow).

## DATA AND CODE AVAILABILITY

Coordinates and structure factors for MUC D1 bound to Cu^2+^ have been deposited in the Protein Data Bank (PDB ID: 7PRL).

## ACKNOWLEDGMENTS

We thank Prof. Byung-Eun Kim and Dr. David Gokhman for helpful suggestions.

## AUTHOR CONTRIBUTIONS

DF, KF, and NR conceptualized and designed research. NR, ADG, KR, YF-S, GJ, NAN, KC, KWR, and DF carried out experiments. YF-S and TI provided methodology and expertise, NR, ADG, KWR, KC, and DF analyzed data. DF wrote the original manuscript draft, and DF, KF, and NR reviewed and edited the manuscript with input from all authors.

## STAR METHODS

### LEAD CONTACT AND MATERIALS AVAILABILITY

Requests for further information and or reagents may be addressed to the Lead Contact, Deborah Fass. The primary plasmids used in this study were deposited to Addgene (identifiers: xxx).

### EXPERIMENTAL MODEL DETAILS

#### Caco-2 cell line

The human colon epithelial cells line Caco-2 was a kind gift from the laboratory of Prof. Benjamin Geiger. Cells were grown in DMEM (Sigma Aldrich) supplemented with 20% fetal bovine serum, L-glutamine, and Pen-Strep (Biological Industries).

### METHOD DETAILS

#### Protein production and purification

The MUC2 D1 coding sequence (NCBI reference sequence NP_002448.4, amino acids 21-389) was cloned into the pCDNA3.1 vector following the signal sequence of the enzyme QSOX1. A His_6_ tag was appended to the carboxy terminus. The plasmid was transiently transfected into HEK293F cells (ThermoFisher) using the PEI Max reagent (Polysciences Inc.) with a 1:3 ratio (w/w) of DNA to PEI at a concentration of 1 million cells per ml. Cells were maintained in FreeStyle 293 medium. To facilitate subsequent crystallization, 5 μM kifunensine (Cayman Chemical) was added at the time of transfection to obtain protein containing high-mannose, EndoH-cleavable glycans. Six days after transfection, cells were removed from the cultures by centrifugation for 10 min at 500g. The culture medium was then further clarified by centrifugation for 30 min at 9500g and filtration through a 0.45 μm pore-size membrane. The D1 assembly was purified from the medium by nickel-nitrilotriacetic acid (Ni-NTA) chromatography. Purified protein was exchanged into 25 mM HEPES, pH 7.5, 250 mM NaCl and concentrated to 10 mg/ml (240 μM).

Protein concentration was determined by diluting an aliquot with 6 M guanidinium chloride in 20 mM sodium phosphate buffer, pH 7 (ε_280nm_ for MUC2 D1 = 43,245 M^−1^ cm^−1^). EndoH (500 units) was added to 400 μl D1 and incubated overnight at room temperature before crystallization.

For experiments aside from crystallization, the MUC2 D1 expression construct was modified to contain a His_6_ tag and tobacco etch virus (TEV) cleavage site following the signal sequence, and the His_6_ tag at the carboxy terminus was eliminated. Protein was produced without kifunensine. After Ni-NTA purification, protein was subjected to cleavage with TEV protease containing a His_6_ tag. The cleaved amino terminal tag and the TEV protease were subsequently removed using Ni-NTA beads. The protein was concentrated in 10 mM MOPS buffer, pH 7.0, 100 mM NaCl. Glycerol was added to 10%, and aliquots were stored at −80 °C until use. An expression plasmid for Muc5b D1 (UniProt E9Q5I3 amino acids 50-426) was prepared with a TEV cleavage site followed by a His_6_ tag at the carboxy terminus. Muc5b D1 was produced and purified as for MUC2 D1, subjected to TEV cleavage, and quantified (ε_280nm_ for Muc5b D1 including the remaining segment of the TEV site = 50,360 M^−1^ cm^−1^).

#### Crystallization and structure solution

The MUC2 D1 assembly containing a carboxy-terminal His_6_ tag was crystallized using the hanging drop method over a well solution containing 30 mM MgCl_2_, 8% polyethylene glycol 3350, 100 mM citrate buffer, pH 5.4, and 10% glycerol. The protein stock solution for crystallization was supplemented with 480 μM copper sulfate. Drops were prepared by mixing 2 μl protein with 1 μl well solution. Data-quality crystals grew slowly over the course of a few months. About 10 min prior to flash freezing, crystals were transferred to a solution containing all components of the well except that the glycerol concentration was increased to 20%. For reduction of Cu^2+^, 5 mM ascorbic acid (Sigma) was included in this soak solution. Data were collected at the European Synchrotron Radiation Facility (ESRF) beamline ID23-1 at 100 K. The wavelength was 14.2 keV (0.8731 Å). Initial phases were provided by molecular replacement using the D1 assembly from the MUC2 amino-terminus cryo-EM structure (Javitt et al., 2020), and the structure model was improved by cycles of rebuilding using Coot (Elmsley et al., 2010) and refinement using Phenix (Adams et al., 2010). No Ramachandran outliers were present in the refined structure model, and 95.5% of the amino acids were in favored regions of Ramachandran space. Though the D1 assembly was previously seen to bind calcium in the cryo-EM structure (Javitt et al., 2020), no calcium was detected in the D1 crystal structure, most likely due to the high concentration of citrate in the crystallization buffer.

#### Isothermal titration calorimetry (ITC)

MUC2 D1 was dialyzed against 20 mM 4-(2-hydroxyethyl)-1-piperazineethanesulfonic acid (HEPES) buffer, pH 7.4, containing 150 mM NaCl and 2 mM, 10 mM, or 30 mM glycine. These conditions were based on another study using glycine as a competitor for copper to enable accurate ITC measurements of high-affinity protein-copper interaction (Trapaidze et al., 2012). After dialysis, protein was diluted to a final concentration of 50 μM. CuSO_4_ was prepared at 500 μM in the same dialysis buffer as the corresponding protein. Experiments were conducted at 25 °C on a Malvern MicroCal PEAQ-ITC system starting with a single 0.4 μl injection of Cu^2+^ solution into protein solution, followed by 18 2-μl injections, with 150 seconds spacing between each injection and continuous stirring at 750 rpm. For each glycine concentration, a blank run titrating the Cu^2+^ into buffer was done and subtracted accordingly. The titration data, including the apparent K_D_, were analyzed using software provided by the manufacturer. The conditional K_D_ values were derived according to previous methods (Trapaidze et al., 2012). The experiments were done in triplicate, and representative measurements are displayed in the figures.

The consistent deviation from a 1:1 stoichiometry in fits to the ITC data (Extended Data Table 2) is not yet explained. X-ray fluorescence counts at the copper edge showed that purified MUC2 D1 was not pre-loaded with Cu^2+^, and measurements after addition of Cu^2+^ suggested stoichiometric binding. Protein concentrations were determined for ITC and EXAFS experiments using the same method and were done with care, such that it is unlikely that large errors in protein concentration determination explain the observations of n = ~0.5.

#### Differential scanning fluorimetry (DSF)

To compare thermal stabilization upon Cu binding by WT MUC2 D1 and mutants, as well as by Muc5b, DSF was performed using a Prometheus NT.48 (NanoTemper) instrument. Unless otherwise indicated, proteins were used at a concentration of 7.5 μM in 50 mM 2- (N-morpholino)ethanesulfonic acid (MES) buffer, pH 5.7, 100 mM NaCl, 0.05% Tween-20. When present, Cu^2+^ was added in the form of CuSO_4_ to a concentration of 7.5 μM (1:1 molar ratio) and ascorbic acid to a concentration of 150 μM. Samples were heated from 25 °C to 95 °C at 1°C/min. Tryptophan and tyrosine fluorescence was monitored by recording the 350/330 nm emission ratio after excitation at 280 nm. DSF experiments done at neutral pH showed the same trends; data are reported for mildly acidic pH because the effect of thermal stabilization by Cu^2+^ was greater and because the D1 Cu^2+^ crystal structure was determined at acidic pH.

#### Microscale thermophoresis (MST)

MUC2 D1 was used at a concentration of 500 nM in 50 mM 3-(*N*-morpholino)propanesulfonic acid (MOPS) buffer, pH 7.0, 100 mM NaCl, 0.05% Tween-20. Protein was incubated with 16 serial dilutions of metal in the range of 50 μM to 1.5 nM for 10 min and centrifuged at 20,000g for 10 min at 4 °C. Tryptophan and tyrosine fluorescence was monitored and recorded using a Monolith LabelFree instrument (NanoTemper). Thermophoresis was analyzed at 70% LED and 20% laser intensity for all samples. MUC2 measurements were done in triplicate. Initial fluorescence was equal in all capillaries before thermophoresis measurements. Muc5b was measured and analyzed in the same manner.

#### X-ray absorption spectroscopy

A solution of 800 μM MUC2 D1 in 50 mM MES buffer, pH 5.7, 100 mM NaCl was incubated with 0.8:1 equivalents of CuSO_4_ via syringe pump to ensure no excess metal was present, then was split into two further samples. One sample was treated with excess ascorbate to ensure the cuprous form of the metalloprotein, and both samples were measured as an aqueous glass in 20% ethylene glycol at 10 K. Cu K-edge (8.9 keV) extended X-ray absorption fine structure (EXAFS) and X-ray absorption edge data were collected at the Stanford Synchrotron Radiation Lightsource on beamline 7–3 with an Si 220 monochromator, in fluorescence mode using a high-count rate Canberra 30-element Ge array detector. A Ni filter and a Soller slit were placed in line with the detector to attenuate the elastic scatter peak. For energy calibration, a Cu foil was placed between the second and third ionization chambers. In the case of the Cu^2+^-loaded sample, scans were taken from fresh surfaces of the sample window and their edges compared to ensure no photoreduction occurred, with a shutter in place in between scan points. Six scans of a buffer blank were averaged and subtracted from the raw data to produce a flat pre-edge and remove any residual Ni fluorescence. Data reduction and background subtraction were done using the EXAFSPAK suite, and the data were inspected for dropouts and glitches before averaging. The EXCURVE program was used for spectral simulations.

#### Cu^1+^ quantification

Samples of 60 μM MUC2 D1 were prepared in 50 mM MES buffer, pH 5.7, 100 mM NaCl. Where indicated, CuSO_4_ was supplied at a concentration of 60 μM and, subsequently, ascorbic acid or quercetin was added to a concentration of 1.2 mM. Following a 5-minute incubation, samples were exchanged into 50 mM MES buffer, pH 5.7, 100 mM NaCl using Zeba™ Spin Desalting Columns 7K MWCO (size exclusion) to remove ascorbic acid or quercetin, when present, and any copper not bound to protein. After desalting, samples were diluted by a factor of three (*i.e*., to 20 μM) into the same buffer containing bicinchoninic acid (BCA) (final BCA concentration 1 mM). Absorbance was measured at 562 nm, and data were converted to Cu^1+^ concentration using an extinction coefficient of 7,900 M^−1^cm^−1^ for the Cu^1+^-(BCA)_2_ complex (Young & Xiao, 2021).

#### Competition Titrations and Data Fitting

UV/vis absorption spectra were recorded in a 1-cm quartz cuvette on an SI Photonics model 420 fiber optic CCD array spectrophotometer located inside a Siemens MBraun glove box under inert nitrogen atmosphere. Cu^1+^ stock solutions were prepared by dissolving Cu(CH_3_CN)_4_PF_6_ in anhydrous acetonitrile and subsequently standardized using the chromophoric ligand bathocuproine disulfonate (BCS). The apparent affinity constants for Cu(I) of WT MUC2 D1 and various mutants were determined by competition titrations with colorimetric ligands (L) by using BCA as L in 50 mM MOPS buffer with 150 mM NaCl, pH 7.0, or ferrozine (Fz) or BCA as L in 50 mM MES buffer with 150 mM NaCl, pH 5.7 (*33*, *34*). Aliquots of 10 mM L stock were titrated into solutions containing 50 μM protein and 25 μM Cu(CH_3_CN)_4_PF_6_ (for L = BCA) or 55 μM protein and 50 μM Cu(CH_3_CN)_4_PF_6_ (for L = Fz) to establish the exchange equilibrium expressed by equations 1 and 2. The concentration of [Cu(BCA)_2_]^3−^ complex was calculated from its absorbance at 562 nm (ε = 7,900 M^−1^cm^−1^). The concentration of [Cu(Fz)_2_]^3−^ complex was calculated from its absorbance at 470 nm (ε = 4,320 M^−1^cm^−1^) (Xiao et al., 2013). Calculations of the other species present in solution were calculated from mass-balance equations. Data plotted as [L] vs. [Cu(L)_2_] were fit to equation 3 using GraphPad Prism software to obtain K_exchange_, where K_exchange_ is designated as K, total [Cu] as M, total [MUC2 D1] as P, [L] as y, and [Cu(L)_2_] as x.

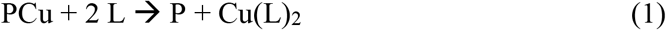

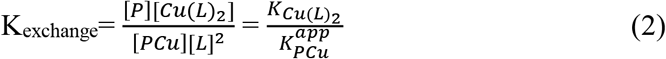

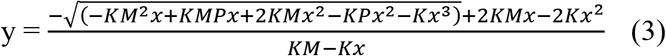

#### Ascorbate Oxidation Assays

UV/vis absorption spectra were recorded in 1-cm quartz cuvettes on a Varian Cary 50 UV/visible spectrophotometer for studies with Cu^2+^. Cu^2+^ stock solution was prepared by dissolving copper chloride (Alfa Aesar) in Nanopure water. Freshly prepared sodium ascorbate (Sigma) was added at a final concentration of 120 μM to buffered 8-μM Cu^2+^-protein solutions. Buffer was 50 mM MOPS, pH 7.0, 150 mM NaCl. Experiments were done in triplicate.

#### Cell Toxicity Assay

Caco-2 cells were passaged in DMEM containing 20% fetal bovine serum (FBS), penicillin/streptomycin, and L-glutamine. Cells were seeded at density of 25,000 cells/well in a 96-well plate, 48 hr prior to experiment. To initiate toxicity, cells were washed once with PBS, and additives for testing were supplied in serum-free medium. Proteins (wild-type or H324A/M326V MUC2 D1) and copper sulfate were supplied at 250 μM final concentrations. After 24 hr, cells were washed with PBS and fixed with 4% paraformaldehyde. After blocking with bovine serum albumin, cells were labelled with anti-E-cadherin antibody ab231303, 1:100 dilution) for 1 hr in room temperature, followed by an Alexa-488 conjugated secondary antibody. Images were taken using a Olympus IX51 microscope equipped with Olympus XM10 camera.

#### COX1 Recovery Assay

Caco-2 cells were passaged in DMEM containing 20% fetal bovine serum (FBS), penicillin/streptomycin, and L-glutamine. For copper starvation, medium was supplemented with 500 μM BCS for 10 days. During passaging under copper starvation conditions, cells were plated in parallel in 6 cm dishes. To initiate recovery, cells were washed once with PBS, and additives for testing were supplied in serum-free medium. After a further 3 days, cells were harvested by trypsinization and pelleted. Cell lysates separate by polyacrylamide gel electrophoresis and analyzed for COX1 levels by western blot (Anti-MTCO1 #ab14705). Bands were quantified using ImageJ software and normalized to the untreated sample for each biological experimental replicate.

## SUPPLEMENTAL FIGURES

**Figure S1.**
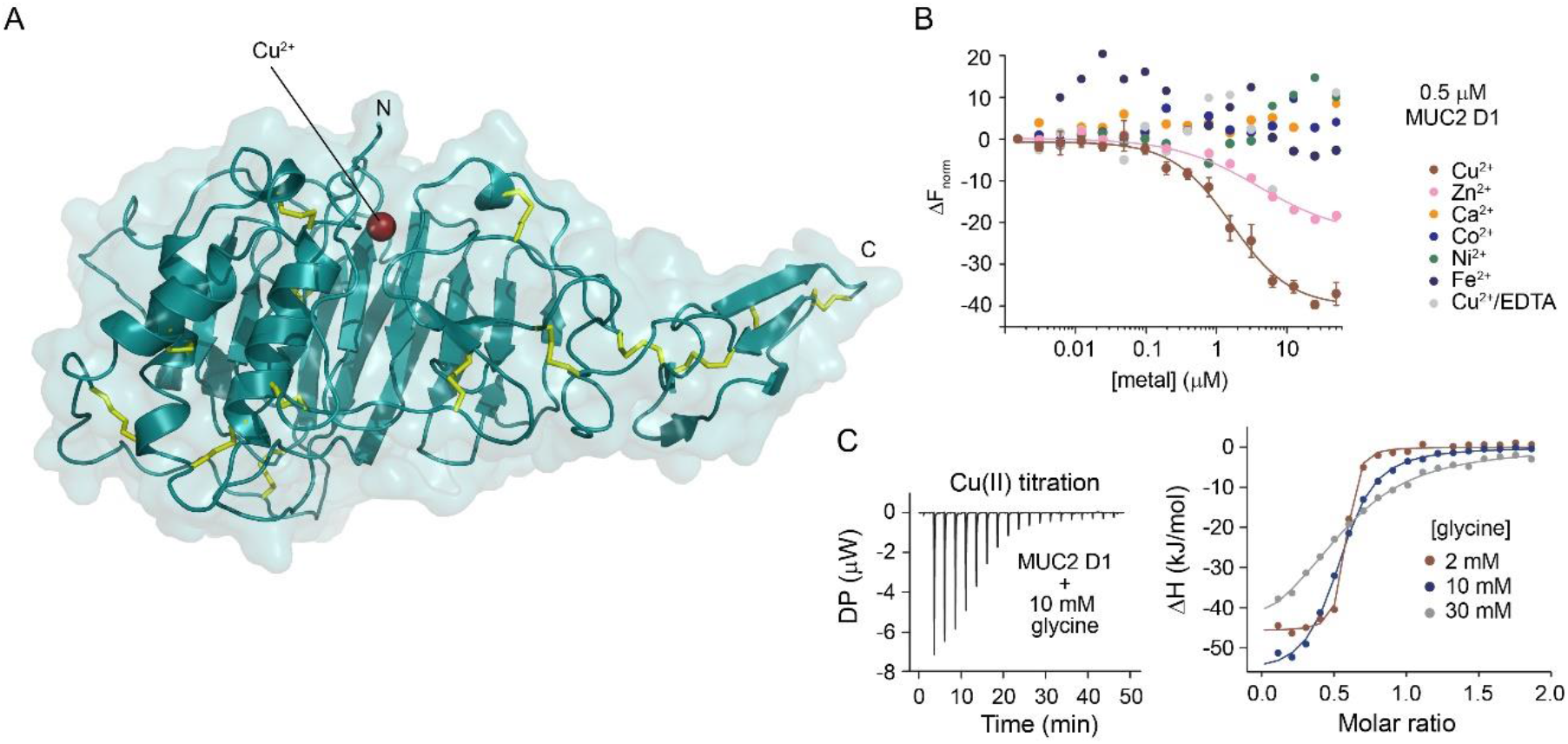
MUC2 specifically binds copper, related to Figure 2. (A) The crystal structure of the Cu^2+^-bound MUC2 D1 assembly is shown rotated 90° around the x-axis compared to Fig. 2A. The Cu^2+^ binding site is near the upper surface of the protein in this view. (B) MST demonstrated binding specificity of MUC2 D1 for Cu^2+^. However, only an upper limit on the binding constant for Cu^2+^ can be determined from this experiment due to the amount of protein required for the measurements (0.5 μM MUC2 D1). (C) ITC measurements in the presence of the competitive ligand glycine. Deviation from a 1:1 Cu^2+^:protein molar ratio is addressed in the Methods.

**Figure S2.**
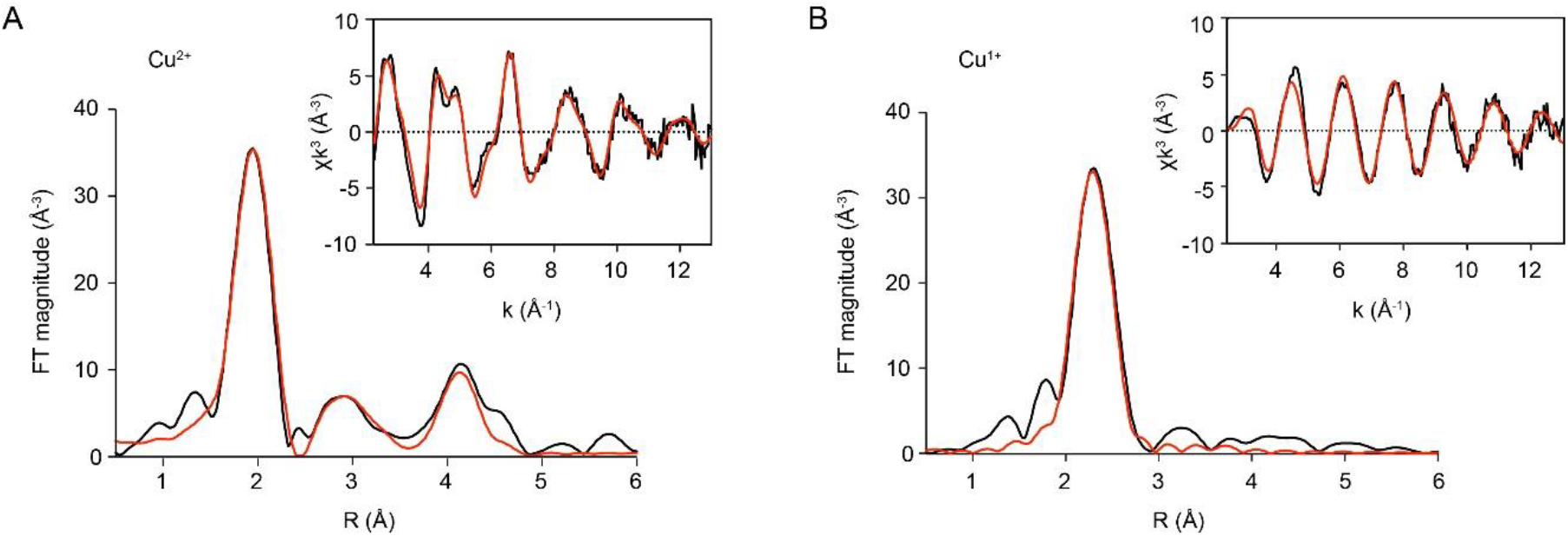
X-ray scattering data for Cu^2+^- and Cu^1+^-bound MUC2 D1, related to Figures 2 and 3. Experimental (black) and simulated (red) Fourier transforms and EXAFS (inset) for Cu^2+^-bound MUC2 D1 and Cu^1+^-bound MUC2 D1 obtained by treating the Cu^2+^-bound protein with ascorbate. (A) An average Cu^2+^–N bond distance of 2.02 Å and Debye Waller (DW) factor of 0.003 Å^2^ were measured for the three His ligands, indicating a stable binding site. The Cu^2+^–N/O bond distance determined by EXAFS, corresponding to the glutamate ligand, was best fit to 1.88 Å. (B) An average Cu^1+^–S bond length of 2.317 Å and DW of 0.011 Å^2^ was calculated for methionine coordination of the reduced copper.

**Figure S3.**
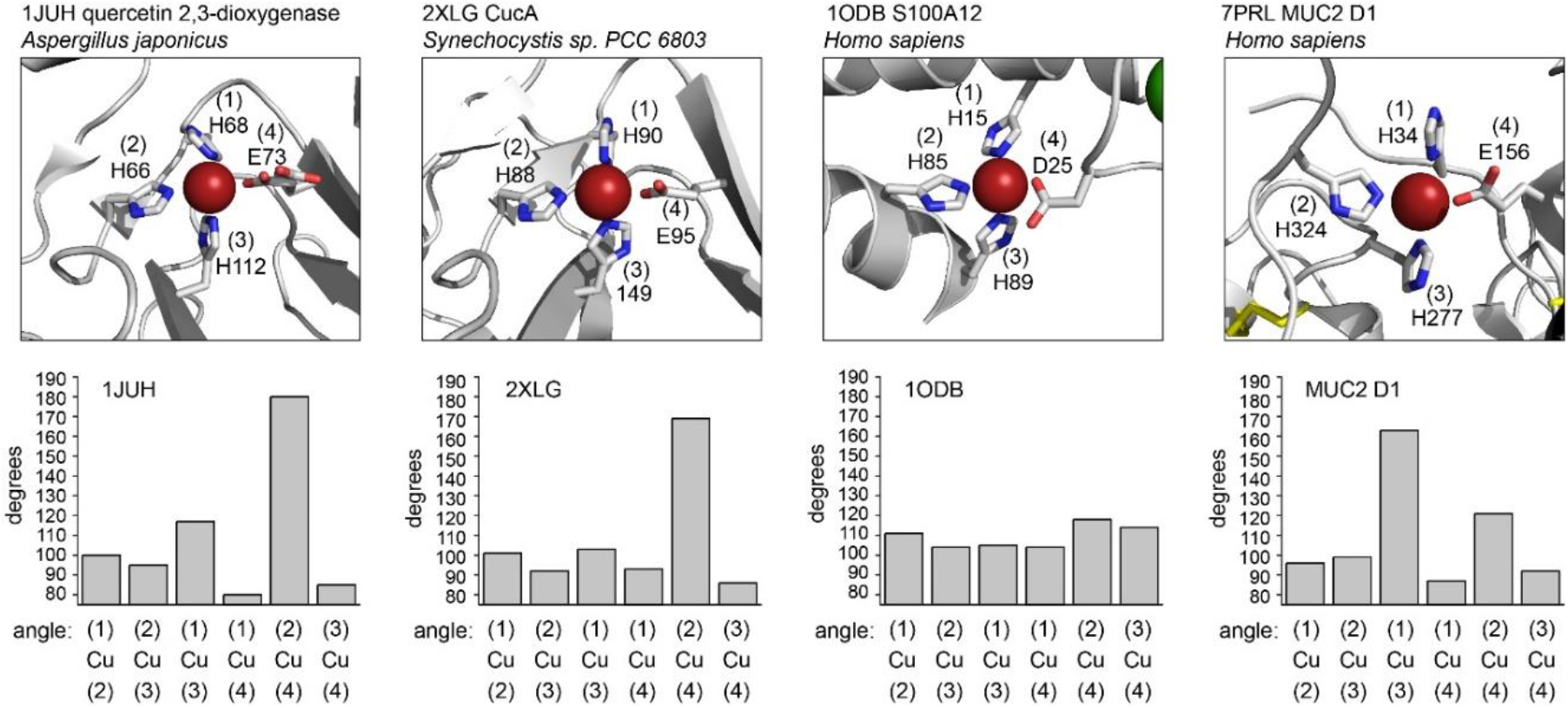
Comparison of copper binding ligand geometries, related to Figure 2. The top panels show examples of protein structures from the protein data bank (PDB) that coordinate copper with similar ligands as the Cu^2+^ site in MUC2 D1. Cu^2+^ coordinating side chains are shown as sticks and numbered according to the amino acid sequence of each protein. In addition, numbers in parentheses designate the coordinating residues counter-clockwise. The lower panels show the angle measured for each pair of coordinating atoms with the copper at the vertex. Quercetin 2,3-dioxygenase (1JUH) and CucA (2XLG) have similar sets of angles, and CucA exhibits some quercetin dioxygenase activity (Tottey et al., 2008). As indicated by all angles being close to 110°, the Cu^2+^ coordination geometry of S100A12 is approximately tetrahedral. The three histidines of MUC2 D1 are arranged roughly as three corners of a square plane (angles ~90°, ~90°, ~180°), while the glutamate is ~120° off the plane. This geometry is different from both 1JUH/2XLG and 1ODB.

**Figure S4.**
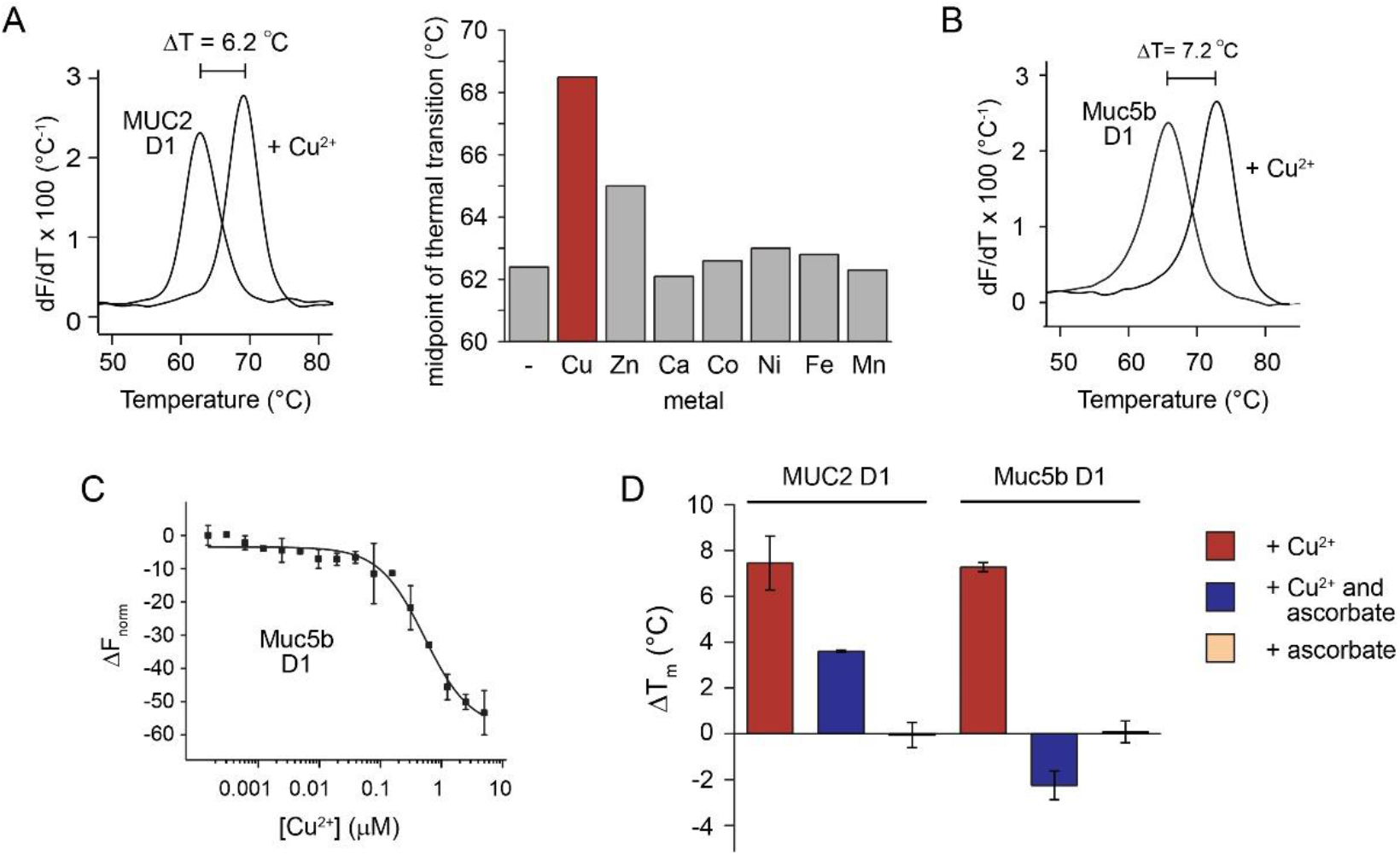
Cu^2+^ increases the thermal stability of MUC2 D1 and Muc5b D1, related to Figures 2 and 3. (A) MUC2 D1 at a concentration of 12 μM was subjected to DSF in the presence of 12 μM of metal. The slope of the fluorescence change is plotted as a function of temperature for the experiment conducted with and without Cu^2+^. A bar graph shows the midpoints of the thermal denaturation transition for all metals measured. Experiments were conducted in 50 mM MES buffer, pH 5.7, 150 mM NaCl, 0.05% Tween-20. Metals were supplied as: Cu, copper sulfate; Zn, zinc chloride; Ca, calcium chloride; Co, cobalt chloride; Ni, nickel sulfate; Fe, iron sulfate; Mn, manganese chloride. Calcium that may have been bound already to the purified protein (Javitt et al., 2020) was not removed prior to the DSF experiment. (B) Cu^2+^ increased the midpoint of thermal denaturation of D1 from the respiratory mucin Muc5b (murine ortholog), measured using DSF. (C) MST demonstrated binding of Cu^2+^ to Muc5b D1, present at 0.5 μM. Experiment was done in duplicate. (D) Unlike Cu^2+^, Cu^1+^, formed by the addition of Cu^2+^ and ascorbate, did not increase the midpoint of thermal denaturation of D1 from Muc5b as measured using DSF.

**Figure S5.**
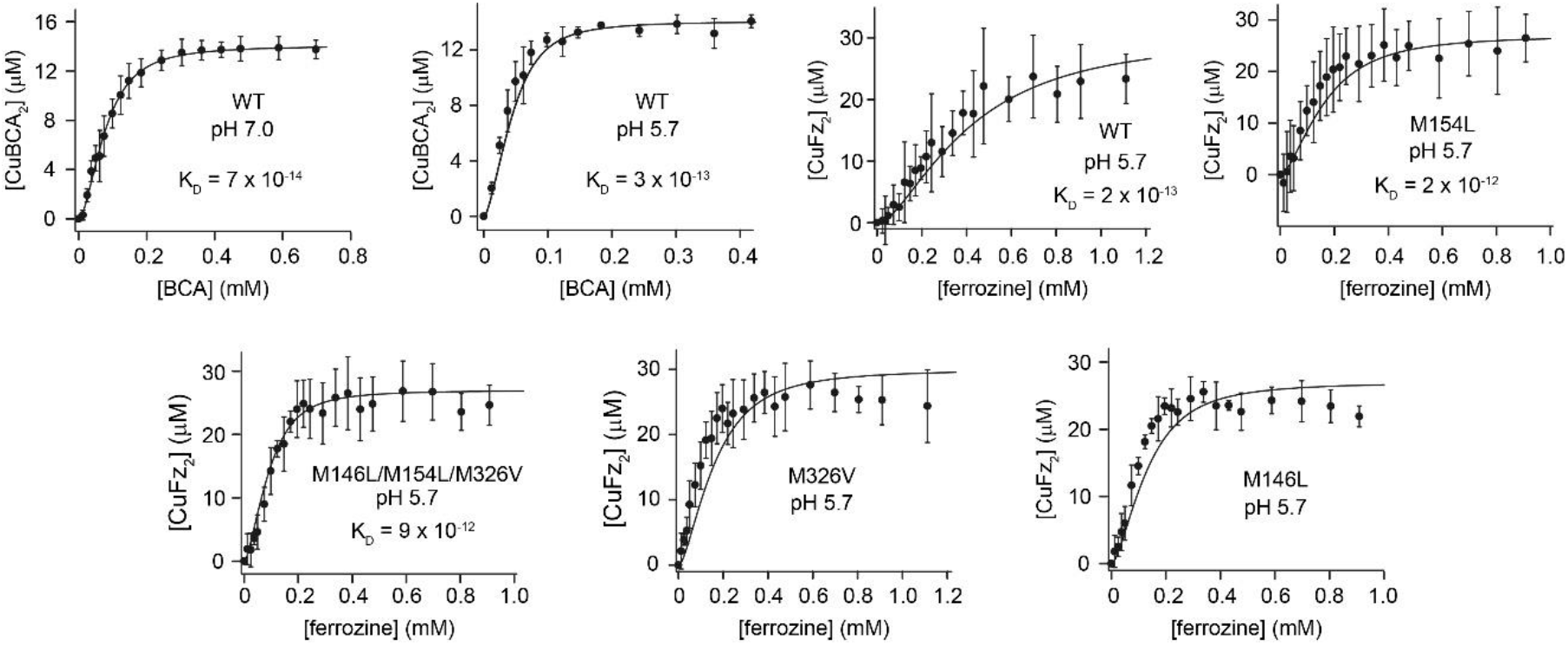
Affinity of MUC2 D1 for Cu^1+^, related to Figure 3. Competitive binding assays of MUC2 D1 with BCA or ferrozine for Cu^1+^ binding under anaerobic conditions. Error bars represent standard deviation from the average of triplicate measurements; the lines represent the best fit to the equilibrium expressed in Eq 3 of the methods section. Dissociation constants (K_D_) obtained from these fits are indicated, except for mutants that did not compete well enough with ferrozine to provide good fits to the data.

**Figure S6.**
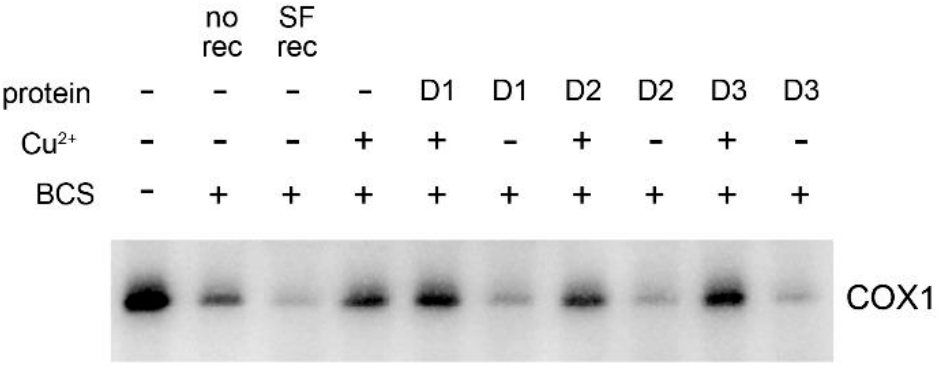
Comparison of COX1 expression after Cu^2+^ addition to copper-starved cells in the presence of MUC2 D1, D2, or D3, related to Figure 5. Caco-2 cells were starved of copper by addition of BCS to the medium where indicated (+) and passaging the cells for 10 days. “No rec” indicates the sample that was then harvested with no recovery. To all other samples, serum-free (SF) medium was supplied with the indicated additives, and cells were harvested 3 days later. When present, proteins were at 5 μM and Cu^2+^ was a 1 μM. Cell lysates were subjected to SDS-PAGE and western blotted for COX1. No appreciable differences were seen between addition of copper together with the copper-binding D1 assembly compared to the non-copper-binding assemblies D2 and D3.

**Table S1.** X-ray crystallographic data collection and refinement statistics

|  | MUC2 D1 + Cu<sup>2+</sup> |
| --- | --- |
| **Data collection** |  |
| Space group | C 2 2 2<sub>1</sub> |
| Cell dimensions |  |
| *a*, *b*, *c* (Å) | 95.90, 141.21, 68.52 |
| $\alpha$, $\beta$, $\gamma$ (°) | 90, 90, 90 |
| Resolution (Å) | 49.17-2.48 (2.57-2.48) |
| *R*<sub>sym</sub> | 0.111 (0.834) |
| *R*<sub>pim</sub> | 0.039 (0.297) |
| *I* / $\sigma$ *I* | 13.4 (2.7) |
| Completeness (%) | 99.2 (99.4) |
| Multiplicity | 9.0 (9.4) |
| **Refinement** |  |
| No. reflections | 16,742 (1,647) |
| *R*<sub>work</sub> / *R*<sub>free</sub> | 0.186/0.260 (0.246/0.331) |
| No. atoms |  |
| Protein | 2750 |
| Ligand/ion | 39 |
| Solvent | 79 |
| *B*-factors |  |
| Overall | 53.5 |
| Protein | 53.3 |
| Ligand/ion | 77.6 |
| Solvent | 49.5 |
| R.m.s. deviations |  |
| Bond lengths (Å) | 0.008 |
| Bond angles (°) | 1.00 |

Values in parentheses are for the highest-resolution shell.

**Table S2.** Thermodynamic parameters obtained by isothermal titration calorimetry for Cu^2+^ binding to MUC2 D1

| Variant | [glycine]<br>(mM) | n | app$\Delta H$<br>(kJ/mol) | app-T$\Delta S$<br>(kJ/mol) | app$\Delta G$<br>(kJ/mol) | appK<sub>d</sub><br>(M) | condK<sub>d</sub><br>(M) |
| --- | --- | --- | --- | --- | --- | --- | --- |
| **WT1**<br>(N31/H34) | 2 | 0.54 | -45.8 | 6.29 | -39.5 | 1.22E-07 | 4.44E-13 |
| **WT1**<br>(N31/H34) | 10 | 0.52 | -57.2 | 23.9 | -33.3 | 1.50E-06 | 2.20E-13 |
| **WT1**<br>(N31/H34) | 30 | 0.57 | -52.7 | 23.8 | -28.9 | 8.59E-06 | 1.40E-13 |
| **WT2**<br>(Y31/R34) | 2 | 0.60 | -57 | 21.5 | -35.5 | 6.09E-07 | 2.21E-12 |
| **WT2**<br>(Y31/R34) | 10 | 0.53 | -71.7 | 40.7 | -31 | 3.74E-06 | 5.48E-13 |
| **WT2**<br>(Y31/R34) | 30 | 0.47 | -97.7 | 71.6 | -26.1 | 2.74E-05 | 4.46E-13 |

WT1 and WT2 are two human MUC2 variants differing at amino acid positions 31 and 34.
<sup>app</sup> = apparent values measured by the ITC software.
<sup>cond</sup> = conditional value calculated according to method described in reference (32).

**Table S3.**
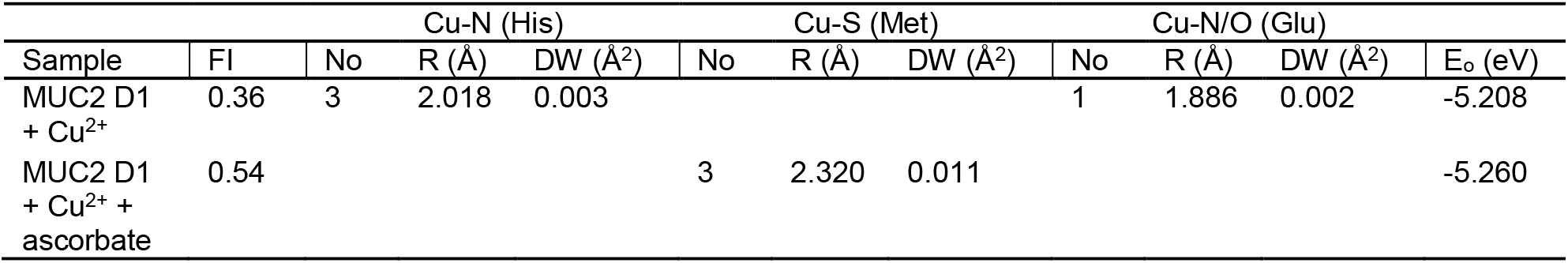
Fitting parameters of Cu K-edge EXAFS spectra

